# Genome-scale metabolic modeling reveals increased reliance on valine catabolism in clinical isolates of *Klebsiella pneumoniae*

**DOI:** 10.1101/2021.09.08.459555

**Authors:** Matthew L Jenior, Mary E Dickenson, Jason A Papin

## Abstract

Infections due to carbapenem-resistant Enterobacteriaceae have recently emerged as one of the most urgent threats to hospitalized patients within the United States and Europe. By far the most common etiological agent of these infections is *Klebsiella pneumoniae*, frequently manifesting in hospital-acquired pneumonia with a mortality rate of ∼50% even with antimicrobial intervention. We performed transcriptomic analysis of data collected from *in vitro* characterization of both laboratory and clinical isolates revealed shifts in expression of multiple master metabolic regulators across isolate types. Metabolism has been previously shown to be an effective target for antibacterial therapy, and GENREs have provided a powerful means to accelerate identification of potential targets *in silico*. Combining these techniques with the transcriptome meta-analysis, we generated context-specific models of metabolism utilizing a well-curated GENRE of *K. pneumoniae* (iYL1228) to identify novel therapeutic targets. Functional metabolic analyses revealed that both composition and metabolic activity of clinical isolate-associated context-specific models significantly differs from laboratory isolate-associated models of the bacterium. Additionally, we identified increased consumption of L-valine in clinical isolate-specific growth simulations. Importantly, valine has been shown to augment macrophage phagocytosis, and this result could be indicative of an immunosuppressive strategy *Klebsiella pneumoniae* evolved for survival during infection. These findings warrant future studies for potential efficacy of valine transaminase inhibition as a target against *K. pneumoniae* infection.

**Importance:** Incidences of infection by *Klebsiella pneumoniae* have grown in frequency to become the leading agents of CRE infection among hospitalized patients in the United States and Europe. Transcriptomic meta-analysis of data collected from both laboratory and clinical isolates indicated significant shifts in expression of key transcription factors related to metabolism. Metabolic network reconstructions have previously proven effective for quickly identifying potential targets *in silico*, therefore we combined these approaches by integrating the transcriptomic data from each isolate type into a well-curated GENRE of *K. pneumoniae* to predict emergent metabolic patterns. Leveraging this systems-biology approach we found discordant patterns of active metabolism between clinical and laboratory isolates, with a striking difference in L-valine catabolism. Exogenous valine is known to increase macrophage phagocytosis, and our results may support immunomodulatory activity in *K. pneumoniae* evolved to avoid host clearance.

## Background

Carbapenem-resistant Enterobacteriaceae (CRE) have emerged as a growing and urgent issue in healthcare facilities around the world, posing a significant threat to public health. Carbapenem antibiotics, currently considered to be the most potent and highly effective class of antimicrobial agents, are often considered a last-resort, reserved specifically for the treatment of severe multidrug-resistant (MDR) bacterial infections^1–3^. This recent surge in CRE-associated infections has been driven primarily by the emergence and dissemination of carbapenemases, a specific type of β-lactamase that has the ability to hydrolyze carbapenems, rendering even carbapenem-class antibiotics ineffective^2^. A large proportion of these CRE-related infections are due to the Gram-negative bacterium *Klebsiella pneumoniae*^2,3^, with over 50% of *K. pneumoniae* infections now being resistant to carbapenems in parts of the Eastern Mediterranean and Europe^3^. As *K. pneumoniae* has been rapidly acquiring antibiotic resistance and rendering almost all available treatments ineffective, the discovery of new treatment strategies for this bacterial pathogen are critical^1,3^.

One strategy that has emerged recently is the targeting of elements of virulence or core metabolism that may be too costly for the organism to accumulate mutations in or diminish the ability to manifest disease^4^. By identifying those characteristics lost during evolution toward sustained laboratory culture, while remaining conserved across infections, it becomes possible to gain insight into important phenotypes that contribute to successful infection. Furthermore, it has been shown that clinical and laboratory isolates of other bacterial pathogens may also be easily differentiated by distinct metabolic capacities^5^. Employing this approach for *K. pneumoniae*, we may highlight “core” metabolic pathways in clinical isolates that may present ideal therapeutic target candidates. Consistent with this strategy, certain elements of metabolism have already been successfully identified as drug targets in bacterial pathogens including other Enterobacteriaceae^6–9^.

Changes in bacterial transcription have been used to assess differences in active metabolism with higher resolution than metabolomics screens, as shifts can be traced to specific pathways and gene products^10^. While RNA-seq has become a relatively standard method for characterizing transcription, technical variability, small sample sizes, and sample heterogeneity still exist and may influence study-specific results^11,12^. Additional differences in data processing criteria also introduce variability into downstream interpretations^13,14^. To account for these factors, meta-analyses of transcriptomic datasets across multiple studies can be performed using a unified curation and analysis pipeline. As such, we assembled 56 publicly available transcriptomes of *K. pneumoniae* isolates from both the laboratory and clinical profiles during growth in similar media conditions at multiple institutions^15–18^. Overall, certain transcriptional patterns varied consistently between clinical and laboratory isolates, and differential expression analysis revealed increased transcription of aminoglycoside degradation and key regulators for histidine utilization among clinical isolates.

To further explore possible targets within infection-associated metabolic pathways, we integrated our transcriptomic meta-analysis with a previously published genome-scale metabolic network reconstruction (GENRE) of *K. pneumoniae* (iYL1228)^19^. GENREs are computational formalisms of the biochemical reactions encoded for in an organism’s genome^20,21^. Previously, GENRE-based growth simulations in other pathogens have successfully highlighted novel enzyme targets which were subsequently validated in the laboratory, effectively accelerating research efforts^8,20,21^. Additionally, GENREs can also be utilized to provide improved context for omics data as the network architecture can reveal additive effects of small changes in activity across interconnected pathways^22^. These network-based analyses enable greater insight into metabolic patterns that correspond with growth under specific conditions, such as during active infection^20,21^. We continued the transcriptomic meta-analysis through integration with metabolic network-based investigation which allowed us to discern novel conserved components of *K. pneumoniae*’s metabolic strategy specific to active infection. Most prominent among these predictions was significantly elevated uptake and utilization of environmental L-valine through the increased activity of an Enterobacteriaceae-specific valine transaminase. This elevated uptake of L-valine was observed across >89% of clinical isolate context-specific models, while nearly entirely absent from laboratory strains, supporting the hypothesis of increased importance for survival *in vivo*. These results also agreed with previous findings that macrophages respond to high concentrations of exogenous valine in order to upregulate phagocytosis and the killing of *K. pneumoniae* during infection^23^. Our study highlights the utility of well curated GENREs integrated with transcriptomic data to accelerate molecular target identification.

## Results

### Transcriptomic data collection

We performed an extensive search of publicly available RNA-Seq datasets on the NCBI Sequence Read Archive^24^ characterizing either laboratory or clinical isolates of *K. pneumoniae*. Data sets were considered for meta-analysis if isolates were grown to exponential phase *in vitro* using LB growth medium at 37°C, at which point transcriptomic samples were collected and sequenced. This standard resulted in 56 total RNA-Seq datasets across four distinct studies; of these datasets, 17 represented laboratory strains^15,16^ and 39 represented clinical isolates^17,18^ (Table S1). The combination of studies from a variety of independent groups help to mitigate concerns due to strain or experimental variation inherent to each individual study^25^.

### Transcriptomic meta-analysis reveals differential expression in metabolic regulators and antibiotic resistance

To first characterize overall transcriptional dissimilarities between isolate types, we performed a differential expression analysis thresholding with an adjusted p-value cutoff of 0.05 and a log_2_ fold change of 2.5 (Figure 1A and Table S2). This analysis identified a total of 19 genes as differentially expressed, 10 that are more associated with clinical isolate and 9 that are more associated with laboratory strains. Among genes with the highest degree of change were a subset of amino acid metabolism regulators, including the hutC transcription factor gene (Figure 2B) and the gcvH glycine cleavage system (Figure 2C), that were both significantly increased in clinical isolates. Specifically, hutC encodes for an established regulator of histidine utilization which has been shown to be a critical colonization factor among certain strains of *K. pneumoniae*^26^. Additionally, transcript for aminoglycoside N-acetyltransferase is significantly overrepresented in clinical isolates (Figure 2D) and encodes for an enzyme that directly mediates the breakdown of aminoglycoside antibiotics. Additionally, a putative stress response protein was expressed more highly in clinical isolates (Figure 1E), and has also been previously associated with increased antibiotic resistance^27^. Alternatively mrkA, a type-3 fimbriae subunit gene, has been shown to facilitate biofilm formation^28^ and whose expression was significantly more associated with laboratory isolates (Figure S1A). Furthermore, genes for multiple ribosomal subunits were highly expressed in laboratory strains (Figures S1B & S1C), suggesting greater optimization for faster growth in culture medium^29^. Cumulatively, these results support the observation that overall transcriptional activity strongly differed between laboratory and clinical isolates in a manner that may impact infection outcomes, and underscored that key metabolic shifts may play a role in these differences.

**Figure 1).**
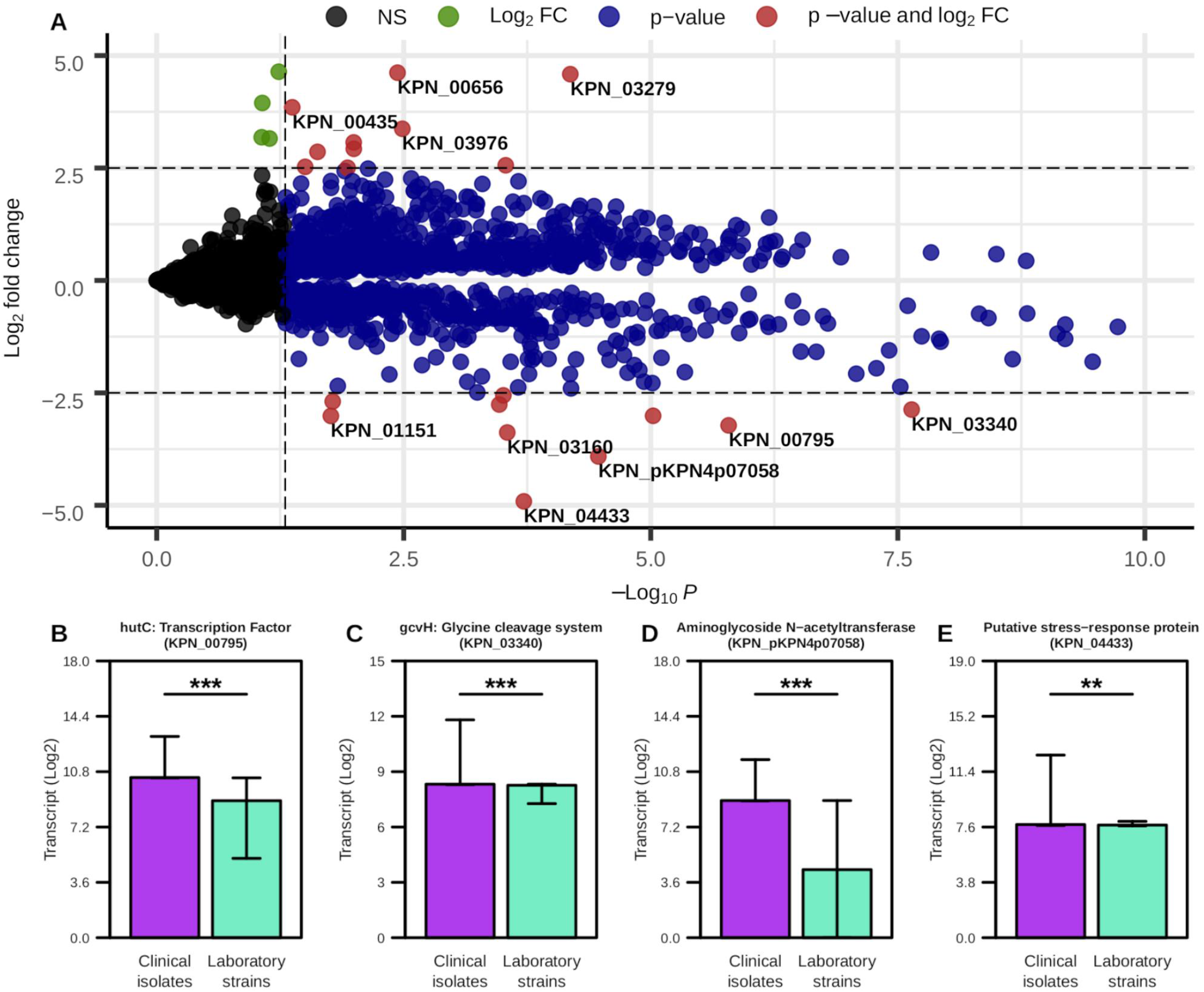
Transcriptional differences between laboratory and clinical isolates of *K. pneumoniae*. **(A)** Differential expression analysis with Log_2_ fold change cut off = 2.5, p-value cut off = 0.05, genes with highest degree of difference are labeled. **(B-E)** Median and interquartile ranges for select genes based on previous analysis. Significant differences determined by Wilcoxon rank-sum test with Benjamini-Hochberg correction (*** *p*-value ≤ 0.001, ** *p*-value ≤ 0.01).

**Figure 2).**
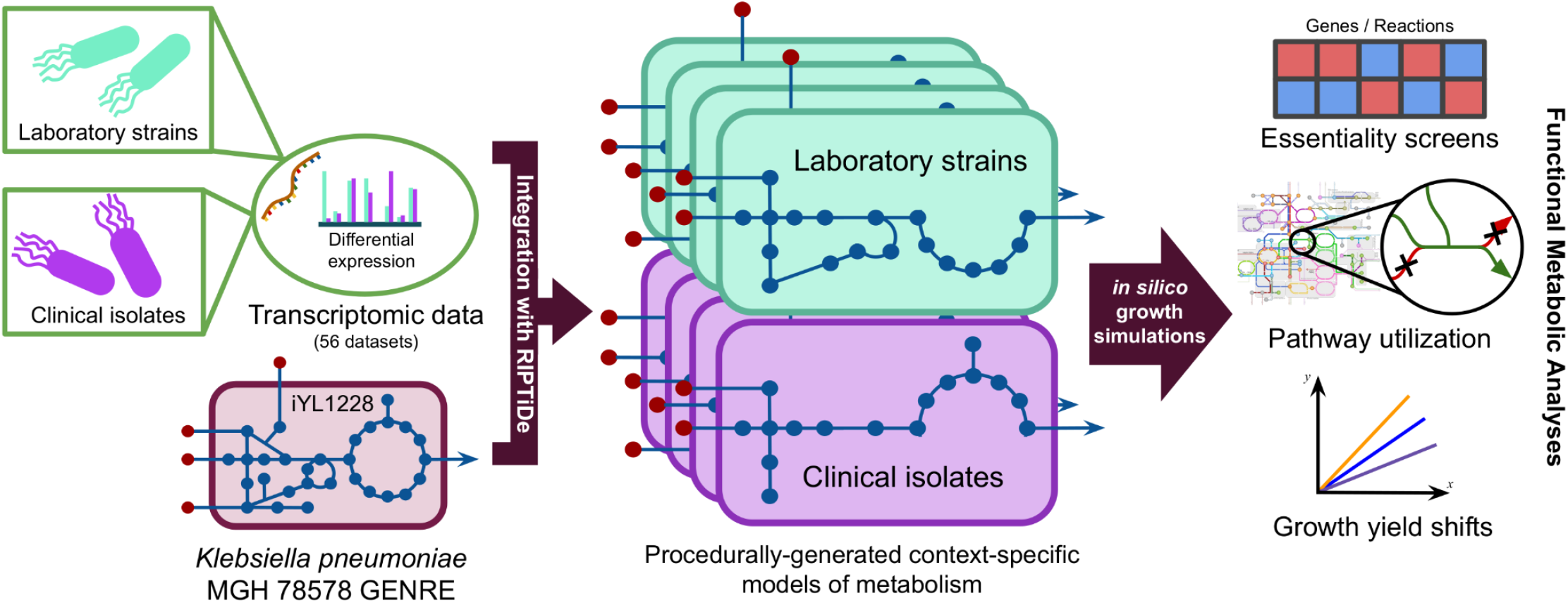
General procedure for generating context-specific models of metabolism from transcriptomic data. All 56 datasets from the transcriptome meta-analysis were used to generate distinct context-specific models of *K. pneumoniae* metabolism.

### Leveraging transcriptomics and GENREs to generate context-specific models of *K. pneumoniae*

Previous studies have shown that GENREs are powerful platforms for transcriptomic data integration, allowing for the capture of greater context surrounding metabolic shifts between varying conditions^22^. We therefore generated context-specific models of metabolism representing either clinical or laboratory *K. pneumoniae* isolates utilizing a recently published method for transcriptome data integration^28^, alongside a well-curated *K. pneumoniae* GENRE (iYL1228)^31^. Briefly, the transcriptomic data integration method identifies the most cost-effective usage of metabolism to achieve growth that best reflects the cell’s investments into transcription and further prunes inactive reactions. Using this approach, we generated unique isolate type-specific models of *K. pneumoniae* metabolism in rich medium for each of the 56 collected transcriptomes, and assessed the emergent differences in active metabolism (Figure 2). The resultant models of context-specific metabolism contained a median of 298 and 302 reactions in laboratory or clinical isolate transcriptome-associated respectively from the total 2262 reactions in the uncontextualized iYL1228 (Table S3). Interestingly, models derived from clinical isolate transcriptomic data were consistently larger than those from laboratory strains, reflecting possible loss of unnecessary metabolite biosynthesis during evolution toward growth in rich *in vitro* culture medium. This data-driven minimization of the possible metabolic solution space more readily reveals critical elements of context-specific metabolism of the organism otherwise not detectable from strictly analyzing the transcriptomic data, and allows for a variety of downstream growth simulations.

### Network topological analysis and essentiality screens highlight valine catabolism as differentially critical in clinical isolates

To begin to assess overall differences in each contextualized model, we performed an analysis of unique subnetwork topology to each isolate type-specific group of models. After subtracting “core” reactions that were present in all 56 models, we were left with a median of 72 and 52 reactions that were unique to either clinical or laboratory-specific models (Table S3). Finally, to focus the analysis on those reactions most shared within each group we further limited the scope of reactions to only those shared by at least 55% of models within each group respectively, revealing 15 differentially active reactions (Figure 3A). Among the most prominent patterns from this analysis were reactions for the import of environmental L-valine in clinical isolate models that were not present in their laboratory counterparts. This finding was interesting as it has been recently discovered that exogenous L-valine promotes increased macrophage phagocytosis *in vivo*, thereby pressuring a lung pathogen to evolve to remove excess valine from the environment^23^.

**Figure 3).**
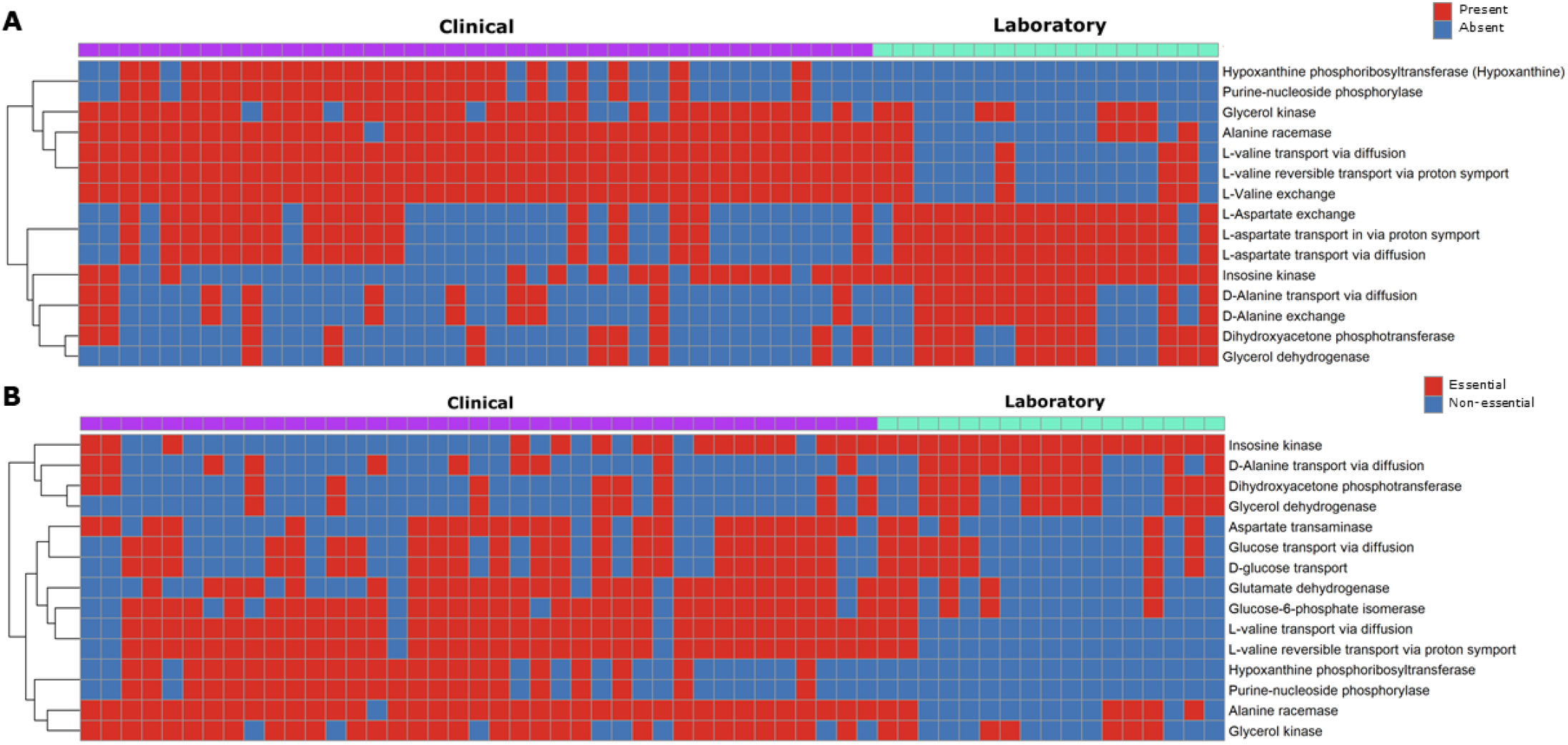
Environmental valine is differentially essential in clinical-isolate context-specific models of *K. pneumoniae*. **(A)** Reaction topology differentially present between the context-specific model groups. **(B)** Differential reaction essentiality between isolate model groups, essentiality was determined through single reaction knockout screen with a cutoff of 1% of the biomass flux. Inclusion in final analysis was determined by cross reference against uncontextualized GENRE and a within-group shared threshold of >55% of models possessing a given feature. Color within the figure area indicates essentiality/presence (red) and non-essentiality/absence (blue), and color on the top margin denotes strain-type of origin for the associated transcriptome with clinical isolate (purple) or laboratory strain (teal).

Next, we sought to identify differentially essential metabolic pathway elements between clinical and laboratory isolates, and ultimately provide a basis for future drug discovery efforts. To accomplish this goal, we performed both single gene and reaction knockout simulations across context-specific models using a threshold of a minimum of 1% of optimal biomass for a gene or reaction to be deemed essential^32^. This functional analysis resulted in a median of 282 and 262 essential reactions in clinical and laboratory associated models, respectively (Table S3). We then cross referenced these results against the uncontextualized iYL1228 to limit potential targets to only those components of metabolism that were environment-specific, and likely not due to strict user-applied constraints, as well as subtracting the “core” essential reactions, resulting in a reduction to a median 72 (clinical) and 52 (laboratory) essential reactions. Then using a similar 55% threshold of shared elements to the previous topology analysis, our combined essentiality screen reported a total of 18 reactions as differentially essential between isolate types (Figure 3B). This analysis indicated that bioconversion of environmental valine is essential in clinical isolates, but not for laboratory strains of *K. pneumoniae*. Of the 18 differentially essential reactions parsed, 3 reactions were directly related to valine metabolism (Figure 3B). These 3 reactions had the highest levels of consistent essentiality among analyzed reactions, being essential for growth in approximately 90% of clinical isolate context-specific models. These results seem to agree with the prior topology-based findings, indicating that valine catabolism may play an important role in the metabolism of *K. pneumoniae* during infection.

### Simulated growth analysis reveals growth advantage and metabolic heterogeneity among clinical *K. pneumoniae* isolates

After observing consistent differences conserved across the metabolic models with individually integrated transcriptomes, we then performed a unified analysis within each group (clinical and laboratory) to incorporate all possible strain-level variation into single models of metabolism for each isolate type. We performed an exhaustive search across biomass flux threshold values to select the level that corresponded with maximum correlation between transcriptome abundances and the network reaction fluxes values. The resultant models account for the variation captured within each isolate type (clinical and laboratory) and the optimal flux distributions that most fit with the transcriptomic investments made by the bacterium. From the corresponding flux distributions, we first measured differences in sampled biomass reaction flux, which is analogous to the growth rate and accounts for biosynthesis of major cellular components (Figure 4A). Strikingly, growth simulations predicted that the clinical isolate-specific model produced biomass at a significantly higher rate than laboratory strains under the same extracellular conditions (*p*-value ≪ 0.001). This observation may be explained by the fact that colonizing a host organism presents substantial environmental pressure and encourages rapid growth to ensure the highest probability of colonization.

**Figure 4).**
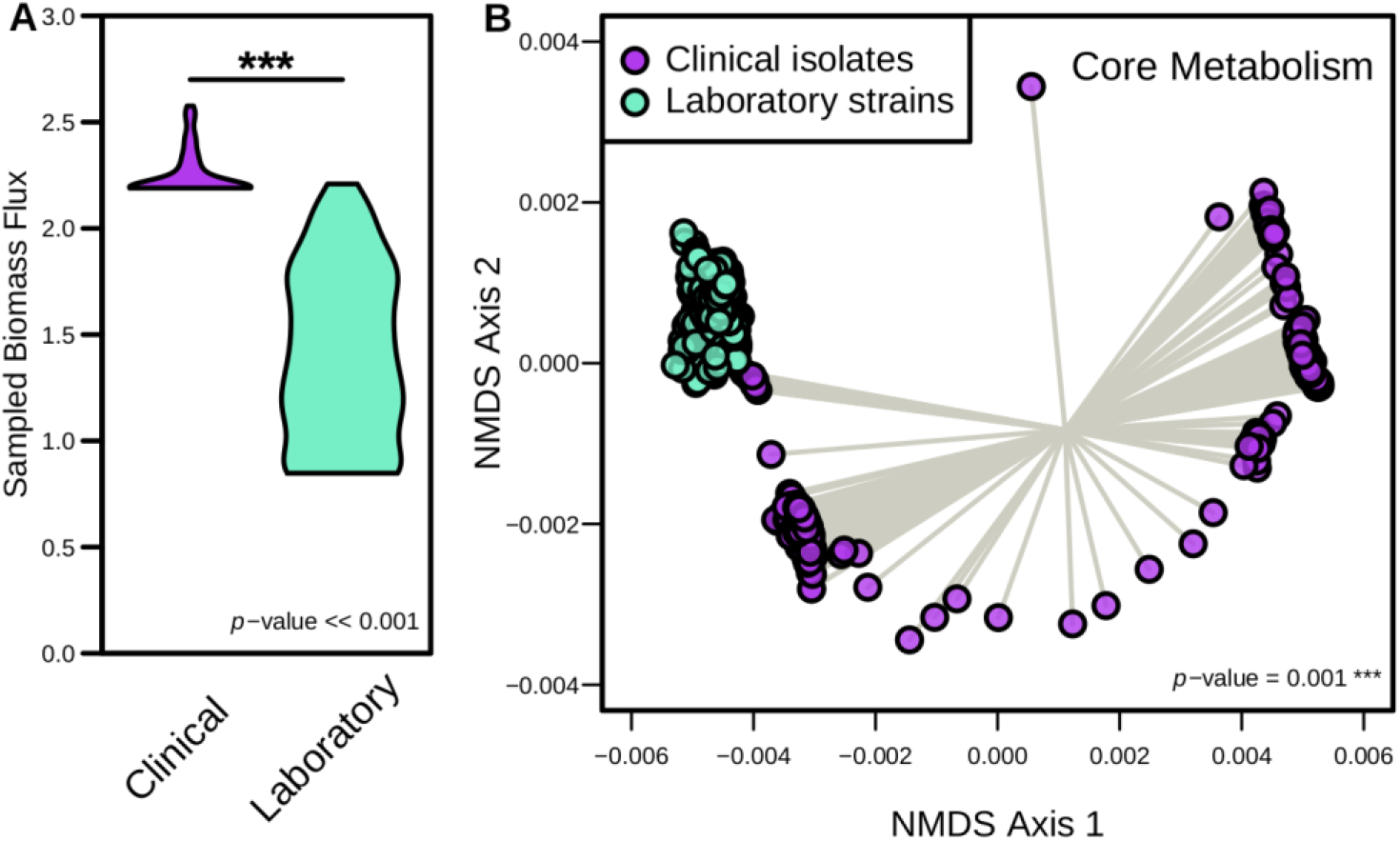
Growth rates and core metabolic activity significantly differ between clinical and laboratory isolate context-specific models. **(A)** Distribution of biomass synthesis reaction flux in each context-specific model, which relates to optimal growth rates. Significant difference determined by Wilcoxon rank-sum test. **(B)** Non-metric multidimensional scaling (NMDS) of Bray-Curtis dissimilarity between sampled context-specific flux distributions from infection and *in vitro* growth. Each point represents the collective activity of all reactions in a context-specific model during optimal growth conditions. Significant difference determined by PERMANOVA.

To then evaluate the degree to which core metabolic activity was altered across isolate types, we focused the analysis of reaction activity on reactions shared across both clinical and laboratory-specific models (excluding the biomass synthesis reaction). Using the reactions flux distributions from this subset of the overall metabolic networks we performed unsupervised machine learning through non-metric multidimensional scaling (NMDS) of Bray-Curtis dissimilarities (Figure 4B), which indeed showed a significant difference in core metabolic activity between the isolate type-specific models (*p*-value = 0.001). This finding was intriguing as it indicated the strain groups are likely adapted for growth in distinct metabolic environments despite simulated growth in the same media conditions. Within-group variation was also significantly greater in clinical isolates (*p*-value < 0.001, Table S4), accurately reflecting the environmental diversity within patients from which they were isolated. Conversely, the variation of core metabolic activity was very low in laboratory-specific growth simulations, potentially suggestive of evolution toward growth in culture medium.

### Dynamic flux analysis further supports valine consumption as important to the metabolic strategy of clinical isolates

Based on our previous results, we investigated differences in L-valine usage between the two isolate-type models during growth. To address this question we utilized dynamic flux balance analysis (dFBA) to simulate L-valine consumption in relation to biomass production over time. Interestingly, in the clinical isolate-specific model L-valine is rapidly depleted from the extracellular environment prior to commencing growth (Figure 5A), while it was consumed at a relatively linear relationship with biomass formation in the laboratory-specific model (Figure 5B). Additionally, although slightly delayed, the maximum growth rate achieved by the clinical isolate model was significantly greater than that of the laboratory isolates (*p*-value ≪ 0.001). Closer investigation revealed the increased L-valine consumption was due to activity of valine transaminase, which mediates the conversion of L-valine to L-glutamate (Figure 5C). This reaction was highly active within the clinical isolate model, but entirely pruned from the laboratory strain-specific model (Figure 5D). Although the gene for valine transaminase (ilvE, KPN_04269) is more highly transcribed in laboratory isolates, activity supporting enzymes in the valine utilization pathway are more active in the clinical isolate-specific model (Figure S2). These results support the hypothesis that the clinical isolates may be more primed to consume environmental valine, despite not being auxotrophic for the amino acid. As previously stated, exogenous L-valine has recently been shown to act in an immunostimulatory manner, causing an upregulation in macrophage phagocytosis in the host immune system^23^. Other studies have additionally shown that glutamate may play an immunosuppressive role, as accumulation of glutamate can lead to limited T-cell function^33^. Combined, these findings suggest clinical isolates of *K. pneumoniae* may have upregulated amino acid catabolism to combat host mechanisms of antibacterial immunity.

**Figure 5).**
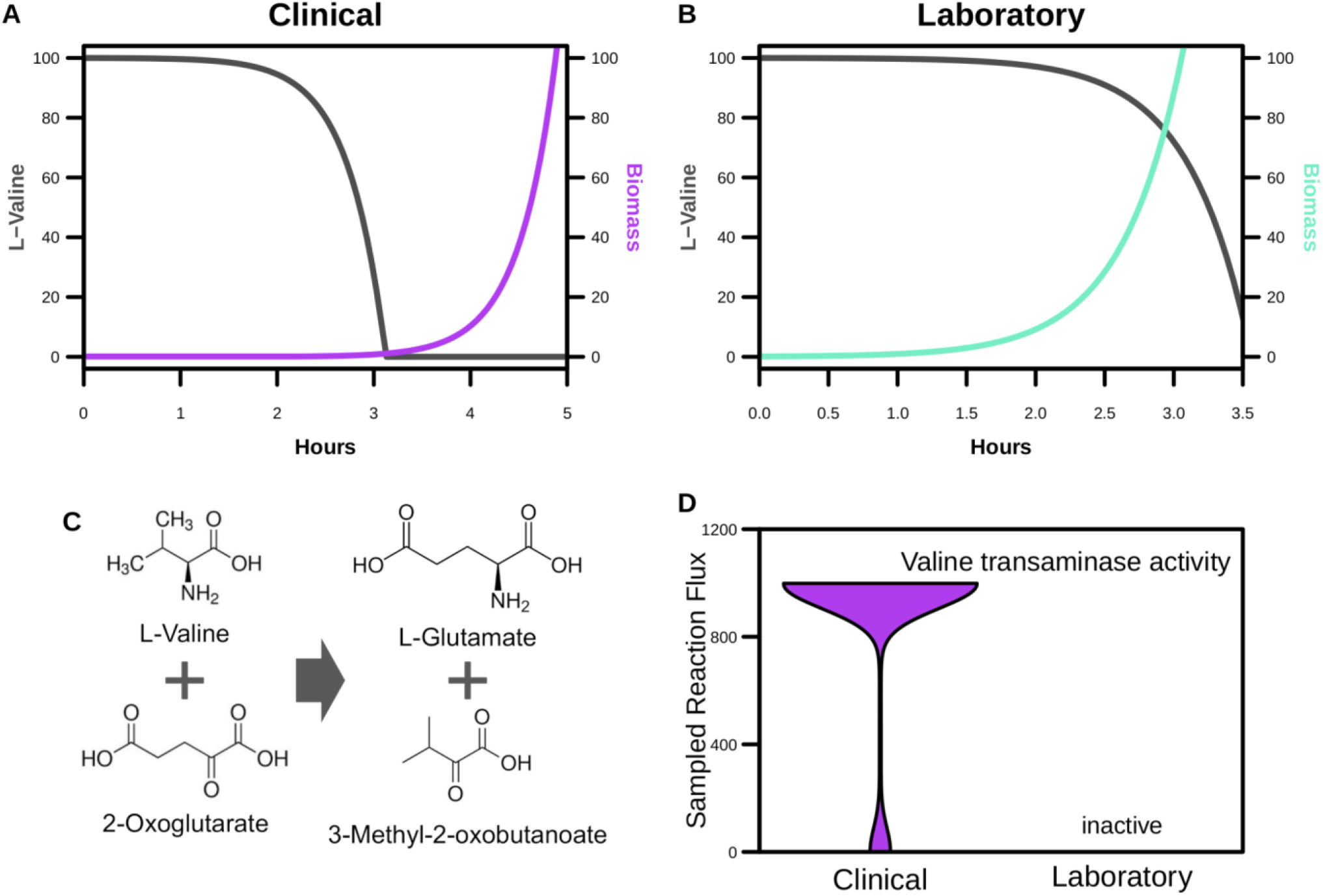
Dynamic flux balance analysis indicates valine consumption prior to exponential growth is more optimal in the clinical isolate-specific model. **(A-B)** dFBA results for each indicated context-specific model. L-valine import flux in mmol/(g h) on the left y-axis and biomass flux in mmol/(g h) on the right y-axis. **(C)** Valine transaminase mediated metabolic reaction responsible for excess L-valine consumption associated with clinical isolate simulated metabolism. **(D)** Valine transaminase-associated reaction sampled flux distributions from each context-specific model. The associated reaction is only present in the clinical isolate-associated metabolic model.

## Discussion

Throughout the past several years, alarmingly increasing numbers of bacterial pathogens have been reported as resistant to antibiotics^34^, emphasizing the need for identification of novel therapeutic options. GENREs have become powerful tools for elucidating the metabolic mechanisms underlying infectious diseases, allowing for the identification and acceleration of novel metabolism-based strategies for treatment^22^. Transcriptomic datasets have additionally been leveraged to further contextualize GENRES to discover essential portions of metabolism for the given cellular states^10^. By targeting elements of metabolism specifically related to life in a host, we may be able to interfere with the ability of the organism to colonize or cause disease. Here, we leveraged computational metabolic modeling of *K. pneumoniae* in combination with transcriptomic meta-analysis to identify unique components of clinical isolate metabolism.

Metabolic modeling results indicated highly distinct patterns of activity within the core metabolism of clinical versus laboratory isolates, highlighting distinct adaptations to their individual environments. Analysis of the models representing clinical isolates point towards the conservation of valine metabolism machinery and the prioritization of early valine catabolism. Additional tracking of the pathways in which valine is metabolized showed that clinical isolates were converting this amino acid into glutamate, which is thought to act as an immunosuppressant. This phenotype may be due to *K. pneumoniae* evolving to sequester valine from the host immune system^23^. The bacteria are then able to convert and excrete the byproducts as glutamate which acts as an immunosuppressant signal. This observation agrees with other studies that have shown the ability to metabolize valine has a clear effect on the fitness of *K. pneumoniae* during active infection^35,36^. Cumulatively, this study points towards the importance of amino acid catabolism for successful host colonization, a functionality that may be conserved among strains more recently isolated from infections.

While this study presents several novel insights into the relationship between the metabolism of *K. pneumoniae* and host factors, some limitations to the analyses are present. While transcriptomic surveys have become relatively standard, there are still potential issues including technical variability and sample heterogeneity which may influence the quality of data in each study^11,12^. Considering these factors, a transcriptomic meta-analysis addresses some of the limiting factors in each component study. We additionally acknowledge that GENREs are not a complete representation for all mechanisms that determine metabolic activity, as they are only built around current reaction annotation data and lack consideration for other levels of regulation^37^. Despite this limitation, the GENRE utilized here was able to accurately predict the metabolic capabilities of *K. pneumoniae* previously^31^, bolstering confidence in the metabolic predictions made here. Furthermore, the discordant relationship with higher valine transaminase transcription in laboratory strains yet lower reaction activity is most likely due to increased transcription of functionally related enzymes in clinical isolates, but inaccuracies in construction of the GENRE may also be a contributing factor that requires additional curation. Despite these considerations, our analyses demonstrate the strength of systems-biology approaches to identify potential metabolic targets against bacterial pathogens.

## Conclusions

Our results indicate that increased valine catabolism is a metabolic phenotype more closely associated with clinical isolates of *K. pneumoniae*. Future studies may build on the targets identified in this study to investigate the role of L-valine in *K. pneumoniae* colonization and virulence or amino acid release and utilization by immune cells. Finally, the methods described here may be applied to other recalcitrant bacterial pathogens in the future as a platform for accelerated drug target discovery.

## Methods

### Transcriptomic Data and Read Curation

All transcriptomic datasets were obtained from the Sequence Read Archive (SRA) in FASTQ format using the SRA Toolkit. Raw reads were quality trimmed using Sickle^38^ to ≥Q25, then strictly mapped to the *K. pneumoniae* MGH 78578 genes (GenBank accession number: CP000647.1) using Bowtie2^39^ and screened for optical/PCR duplicate reads with Picard MarkDuplicates^40^. Mapping files were converted to human-readable format using SAMtools^41^, and transcript abundances were normalized to both read and target gene lengths then evenly subsampled for equal comparison across conditions.

### GENRE-based analyses

The GENRE of *Klebsiella pneumoniae* strain MGH 78578, iYL1228^31^, was obtained from the BiGG Model database^42^ on 5/21/20. Flux analyses performed in this study utilized cobrapy (v0.22.1)^43^. Growth simulations were performed using a previously published rich medium *in silico* formulation^44^. Gene and reaction essentiality screens were both performed with a minimum objective flux threshold of 1.0% of the optimal value. Replicate GENRE transcriptome integration was performed with RIPTiDe (v3.2.3)^30^ with 0.75 minimum objective flux fraction. Maximum fit RIPTiDe analysis was performed with all transcriptome replicates on the default settings.

### Statistical analysis

Statistical analyses were performed in R (v3.2.0). Ordination analysis was accomplished using the vegan package (v2.5.7)^45^.

## Acknowledgments

The authors declare no conflicts of interests. The authors would like to acknowledge Drs. Jhansi Leslie and Kim Walker for many helpful conversations on *Klebsiella* virulence factors, gene regulation, and metabolism. This work was supported by funding from The U.S. National Institutes of Health award R01AI154242 to JP as well as a pilot grant from the UVA Trans-University Microbiome Initiative to MJ.

## Supplementary Materials

**Figure S1).**
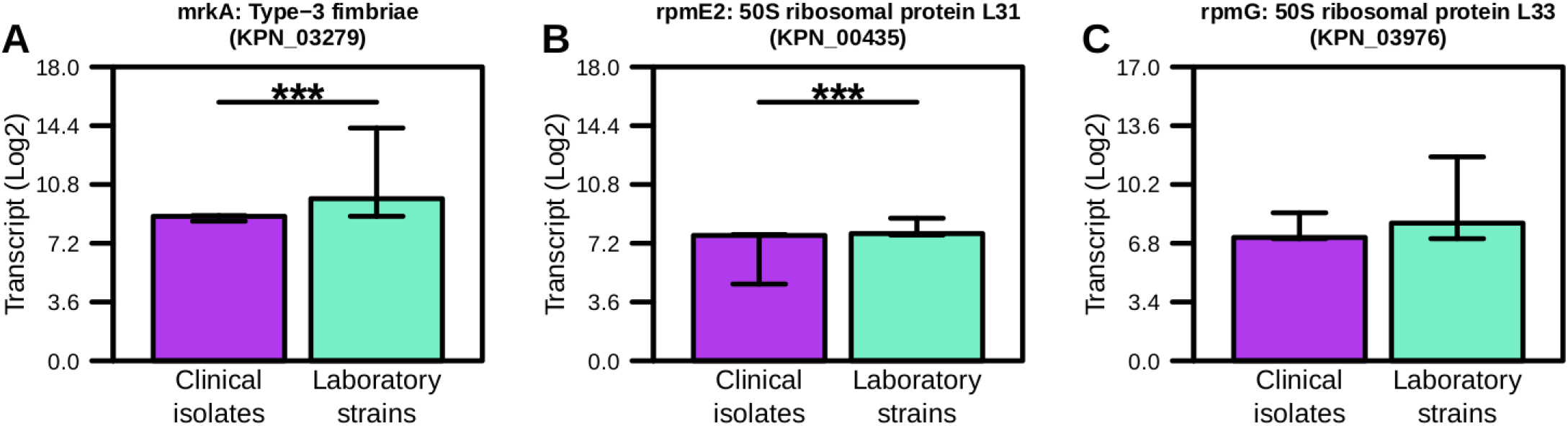
Additional differentially expressed genes from Figure 1. Median and interquartile ranges from pooled analysis between clinical and laboratory isolate transcriptomes. Significant differences determined by Wilcoxon rank-sum test with Benjamini-Hochberg correction (*** *p*-value ≤ 0.001).

**Figure S2).**
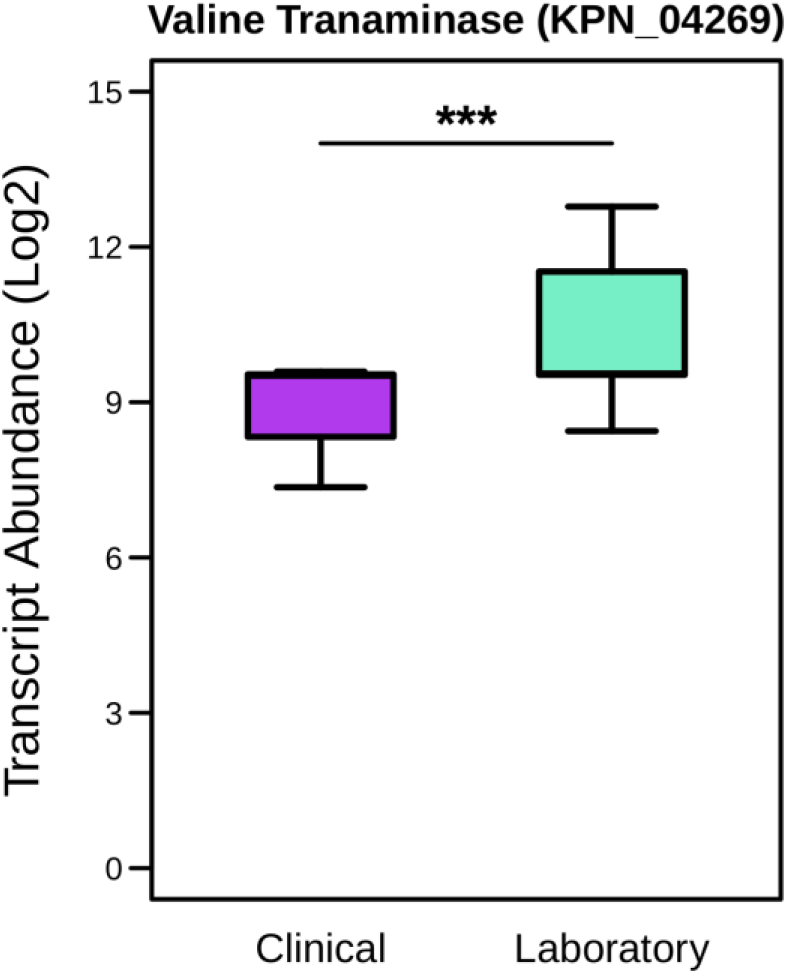
Valine transaminase differential expression. Differential expression analysis across all analyzed transcriptomes. Significant difference determined by Wilcoxon rank-sum test (*** *p*-value ≤ 0.001).

**Table S1** | ***K. pneumoniae* transcriptomic dataset metadata**

**Table S2** | **Differential expression analysis summary statistics**

**Table S3** | **Complete gene and reaction essentiality results**

**Table S4** | **Flux analysis summary table**

